# Integrating over uncertainty in spatial scale of response within multispecies occupancy models yields more accurate assessments of community composition

**DOI:** 10.1101/143669

**Authors:** Luke Owen Frishkoff, D. Luke Mahler, Marie-Josée Fortin

## Abstract

1. Species abundance and community composition are affected not only by the local environment, but also by broader landscape and regional context. Yet determining the spatial scale at which landscapes affect species remains a persistent challenge that hinders ecologists’ abilities to understand how environmental gradients influence species presence and shape entire communities, especially in the face of data deficient species and imperfect species detection.
2. Here we present a Bayesian framework that allows uncertainty surrounding the ‘true’ spatial scale of species’ responses (*i.e.,* changes in presence/absence) to be integrated directly into a community hierarchical model.
3. This scale selecting multi-species occupancy model (ssMSOM) estimates the scale of response, and shows high accuracy and correct type I error rates across a broad range of simulation conditions. In contrast, ensembles of single species GLMs frequently fail to detect the correct spatial scale of response, and are often falsely confident in favoring the incorrect spatial scale, especially as species’ detection probabilities deviate from perfect.
4. Integrating spatial scale selection directly into hierarchical community models provides a means of formally testing hypotheses regarding spatial scales of response, and more accurately determining the environmental drivers that shape communities.

## Introduction

Features of the landscape beyond the local scale often affect the processes that give rise to patterns of community composition (Wiens 1989; Levin 1992; Kneitel & Chase 2004; Dray *et al.* 2012; Fortin *et al.* 2012; McGarigal *et al.* 2016). As a result, ecologists have sought to quantify what landscape features, in what contexts, and at what spatial scales explain the presence and abundance of species. Yet determining how species respond to the landscape has been challenging, in part because the relevant spatial scale(s) at which environmental conditions affect species and communities are rarely known *a priori.* This difficulty has led to uncertainty regarding the conclusions of many landscape level studies (Jackson & Fahrig 2015). The development of statistical methods that more robustly incorporating scales of responses within the statistical analysis of communities (Borcard & Legendre 2002; Jombart *et al.* 2009; Matthiopoulos *et al.* 2011; Dray *et al.* 2012; Warton *et al.* 2015; Ovaskainen *et al.* 2016, 2017), and more accurately convey uncertainty regarding these scales (Chandler & Hepinstall-Cymerman 2016), have the potential to accelerate basic and applied ecological research.

When considering landscape level effects on species presence, abundance, or biomass, two properties of the species are generally of interest. First, at what spatial scale does the species respond to the environment (Desrochers *et al.* 2010), and second, how do they respond (positively or negatively)? The most commonly used approach for determining spatial scale of response *(i.e.,* the spatial context, spatial contingency; Fortin *et al.* 2012) quantifies the average environmental value within buffers of various radii (Holland *et al.* 2004; Weaver *et al.* 2012; Zuckerberg *et al.* 2012; McGarigal *et al.* 2016), and then repeats a statistical analysis using the environmental covariate at each spatial scale (Figure 1). For each species in turn, or for some community-level index like species richness or diversity *(e.g.,* Shannon Index), the most likely spatial scale (as quantified by AICc, correlation coefficient, or slope parameter value) is selected to represent the best match of a species’ response to landscape heterogeneity.

**Figure 1.**
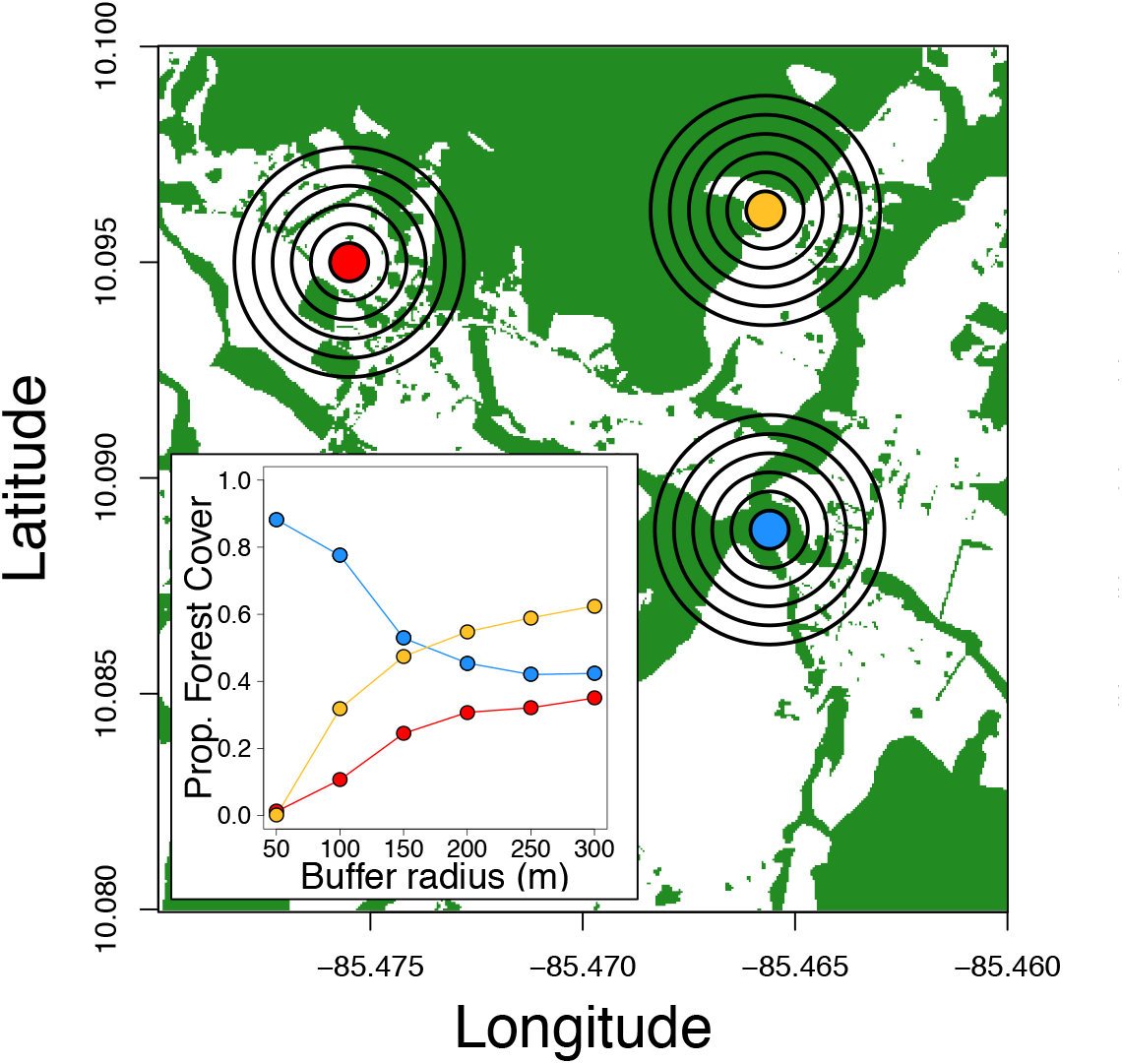
Map of empirical 5m-resolution subsection of forest cover landscape in northwestern Costa Rica with three fictitious sampling points that break the correlation structure between local and broader landscape forest cover. Colored points represent 50m-point count radii, and each successive buffer represents an increase in radius of 50m. Only 50-300m radii are shown, though for simulation analyses radii extended to 1500m.

This multi-scale analysis approach has been successful in elucidating species and community responses (McGarigal *et al.* 2016). By considering landscapes as a whole it has helped quantify the benefits of small forest fragments to biological communities, and related ecosystem services (Karp *et al.* 2013; Mendenhall *et al.* 2016). More generally it has highlighted that species respond to different environmental conditions at different spatial scales, and that species distribution models possess greater predictive power when these multiple scales are directly incorporated (Desrochers *et al.* 2010; Weaver *et al.* 2012). However, the current multi-scale approach does present a number of problems related to estimating the spatial scale of response, exacerbating uncertainty by treating species individually rather than the community as an integrated whole, and ignoring issues with species detectability. All of these will inflate error in estimating the true spatial scale of response, and quantifying how species respond to the environment.

First, single species model comparison approaches that select a single best model typically neither quantify nor integrate over uncertainty regarding scale selection. This means that other parameters may be biased if the ‘most likely’ scale is not the true scale. Relatedly, the set of scales analyzed is often quite small (Desrochers *et al.* 2010; Jackson & Fahrig 2015), and as a result is unlikely to even include the true spatial scale. Meta-analysis has shown that the most likely spatial scale is often at one of the extremes of those analyzed—suggesting that the true spatial scale is even more extreme (Jackson & Fahrig 2015). Recently, Chandler & Hepinstall-Cymerman (2016) proposed a modeling approach that internalizes spatial scale estimation within a single species model by using smoothing kernels to average landscape variables around focal sites. This single-species model addresses these two problems by maximizing likelihood over the spatial scale parameter, and also allows confidence intervals to be calculated around it. Further, because spatial scale is a continuous (albeit bounded) parameter, it eliminates the problem of not including the true spatial scale among the scales assessed (provided an appropriate range of scales is investigated).

The second major problem with the standard approach is that it makes assessments of multiple species in a species-by-species fashion. In the study of entire communities, however, this approach is often inadequate, because it ignores rare species (which are both typically of greatest conservation concern, and may be trophically influential). This problem is especially true in tropical communities, which are particularly under threat from land-use change, and where species rarity is common (MacArthur 1969; Hubbell 2001). Further, a species by species approach is prone to estimation bias and loss of power (Ovaskainen & Soininnen 2011; Banks-Leite *et al.* 2014). Hierarchical joint community models have been proposed to move beyond piecemeal assessments (Ovaskainen & Soininnen 2011; Warton *et al.* 2015; Ovaskainen *et al.* 2017). By assuming that species parameters come from common distributions, overall community error is minimized and rare species can be included in analyses.

Finally, imperfect detection of species is a problem for animal communities generally, especially when species traits, site characteristics, or the time or conditions of observation influence detectability. Multi-species occupancy models (MSOMs) are a commonly adopted solution to account for imperfect and variable detection within communities (Iknayan *et al.* 2013). MSOMs are typically implemented in a Bayesian framework (relying on MCMC to overcome challenges in maximizing likelihoods when numerous random effects exist). Yet, model comparison and selection is still difficult to implement with Bayesian models (e.g., Hooten & Hobbs, 2015; but see Lele *et al.* 2007). Together this means that comparing models across a large range of spatial scales would be both time extremely time consuming (because MCMCs are relatively slow), and non-trivial to implement. Because MSOMs are not easily amenable to adequate testing at multiple spatial scales, the scale of response has generally not been incorporated into community analyses that incorporate imperfect detection. Consequently, their power has not been sufficiently directed to understand how landscape features structure community-level processes.

Fortunately scale selection can be integrated directly into MSOMs by establishing a parameter that allows the spatial scale at which species respond to be estimated (e.g., Chandler & Hepinstall-Cymerman, 2016). When extended to the community as a whole, incorporating scale selection into the model should result in the appropriate spatial scale(s) being estimated directly from the data within a single model run, and also ensure that other parameter estimates are not biased because they were analyzed at the wrong spatial scale. One of us (LOF) recently developed the foundations of the approach presented here in an attempt to internalize scale selection within hierarchical multispecies models to overcome uncertainty about what spatial scale to analyze data for two specific empirical studies (Frank et al. In Press; Karp et al, In review). However, this technique was not fully developed in those works, and it remains unclear whether this approach has correct type I error rates, and whether it does indeed increase power and accuracy over more traditional approaches. Here we fully describe, and demonstrate the use of, a multi spatial scale selection multi-species occupancy model (hereafter; ssMSOM), and test its performance in estimating both spatial scale of response, and species’ strengths of response to the environment *(i.e.,* a landscape-level covariate). We compare this approach to the standard method of analysis: a series of species-by-species GLMs. While the approach here is demonstrated with a simple single season occupancy model for multiple species, the internalized scale selection is generalizable. For example, it could be directly integrated into abundance models, or combined with flexible hierarchical modeling approaches to query population dynamics through times, or the effects of species traits, phylogenetic relatedness, or interspecific competition on community structure (Yackulic *et al.* 2014; Frishkoff *et al.* 2017).

## Methods

### Model overview

The scale selecting multi-species occupancy model (ssMSOM) estimates parameters related to occupancy and detection probabilities in communities containing multiple species (indexed by i), across multiple sites (*j*), with multiple site visits (*k*). The observed detection histories (*Y_i,j,k_*) are assumed to derive from unobserved (latent) occupancy states (*Z_i,j_,* where *Z_i,j_* = 1 for presence, and 0 for absence) and detection probabilities determined by species, and site (*P_i,j_*; visit based variability in detection is ignored for simulations and models, but could be incorporated if desired). Specifically:

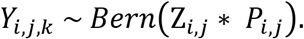

The occupancy state (*Z_i,j_*) in turn was assumed to come from some underlying occupancy probability according to:

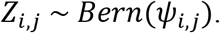

The detection process was modeled according to:

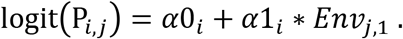

And the occupancy probability (*ψ_i,j_*) according to:

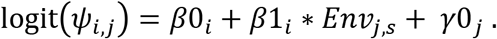

Here *Env_j,s_* is a site-by-scale matrix of some landscape level environmental variable (centered and scaled within each column). All parameters in the *α* and *β* groups are estimated for each species, with species terms drawn from a normal distribution of mean (*μ*) and variance (*σ*^2^) estimated from the data. *γ* terms were random intercepts for each site (variance estimated from data around a mean of 0) designed to incorporate consistent differences in occupancy probabilities in all species across sites that are not accounted for by *Env*. The indexing value *s* (representing columns of the *Env_j,s_* matrix) spans multiple spatial scales, and the parameter value of s that best fits the data is estimated from the model. Because models are implemented using MCMC, this process results in a posterior distribution for values of *s*, which fully integrates over the uncertainly regarding the proper spatial scale, and which further can be used to select the most appropriate spatial scale (*e.g.,* posterior mean or mode) or an interval of spatial scales that well describe the data. This formulation is conceptually similar to generalized linear models that integrate over phylogenetic uncertainty in tree topology (de Villemereuil *et al.* 2012). For simplicity, the environmental effect of a species’ detection probability is assumed to come from the environmental conditions at the finest (*i.e.,* most local) spatial scale. This assumption could be relaxed if there is reason to expect that more distant environmental conditions somehow affect detection probability.

For demonstration and testing purposes, we here assume that a single environmental gradient affects community composition. However, the ssMSOM could be generalized to include multiple environmental conditions (multiple site-by-scale environment matrixes, e.g., *Env1_j,s_ Env2_j,t_ Env3_j,u_*, etc.), each affecting communities at different spatial scales (with scale parameters *s, t, u,* etc., each independently estimated from the data).

### Simulation conditions

In order to test the performance of the ssMSOM, we simulated communities, using an *Env* matrix based on empirical landscape forest cover. Spatial forest cover data for simulation and analysis came from northwestern Costa Rica, used as part of a study of how local and landscape level habitat conversion affects community composition (Karp et. al. In review). In that study, sites were selected to ensure that local forest cover varied independently from landscape level forest cover. To measure surrounding forest cover, all tree cover within 1.5km of sites was classified using high-resolution Google Earth images obtained from 2013-2016. The resulting 5m-resolution tree cover map was verified based on ground-truthed data collected in the field. For analysis, site level forest cover proportion was calculated in radii from 50m to 1500m, in 50m increments, resulting in an *Env* matrix with 30 columns. For the ssMSOM it would be most appealing to use the smallest increments possible to generate the largest number of spatial scales possible, since using few spatial scales makes it likely that the true scale will not be among the analyzed set. We settled on 50m increments because of a balance of computational efficiency in model runs, and increments that approximate a continuous stretch from our smallest to largest spatial scale.

To test the performance of ssMSOM under a variety of conditions, we simulated 120 communities (four at each of the 30 spatial scales), each with 16 species (N_sp_), across 50 sites (N_site_), with three site visits per site (N_visit_). All species parameters were drawn from normal distributions, generating diversity in species’ commonness (simulations included both common and rare species), and species’ responses to tree cover (some responded negatively and others responded positively to ‘deforestation’). This diversity of overall commonness and responses to the environment mimics patterns observed in many empirical systems. We repeated these simulations under five alternative detection scenarios:

1. Perfect detection, where the probability of detecting a species at a site if the site is occupied is 1.
2. High detection probability: average detectability equals ~ 0.5.
3. Low detection probability: average detectability equals ~ 0.25.
4. Low detection with detection affected by local environment: average detectability equals ~ 0.25, but rising to ~0.5 under high local values of *Env* and dropping to ~0.1 under low local values of *Env*.
5. Low detection with species-specific variation in detectability by local environment: Same as 4, but some species increase in detectability with increasing values of local *Env,* while others decrease in detectability.

For an overview of all simulation parameters see Table 1.

**Table 1:** Simulation conditions of communities.

| Component | Parameter | Detection Regime |  |  |  |  |
| --- | --- | --- | --- | --- | --- | --- |
|  |  | Perfect | High | Low | Low-Env | Low-Env Variable |
| Detection | $\mu_{\alpha 0}$ | 100 | 0 | -1 | -1 | -1 |
| | $\sigma_{\alpha 0}$ | 0 | 1 | 1 | 1 | 1 |
| | $\mu_{\alpha 1}$ | 0 | 0 | 0 | 1 | 1 |
| | $\sigma_{\alpha 1}$ | 0 | 0 | 0 | 0 | 1 |
| Occupancy | $\mu_{\beta 0}$ | -1 | -1 | -1 | -1 | -1 |
| | $\sigma_{\beta 0}$ | 1 | 1 | 1 | 1 | 1 |
| | $\mu_{\beta 1}$ | 0.5 | 0.5 | 0.5 | 0.5 | 0.5 |
| | $\sigma_{\beta 1}$ | 1 | 1 | 1 | 1 | 1 |
| | $\sigma_{\gamma 0}$ | 0.1 | 0.1 | 0.1 | 0.1 | 0.1 |
|  | s | variable | variable | variable | variable | variable |
| Sample Size | $N_{sp}$ | 16 | 16 | 16 | 16 | 16 |
| | $N_{site}$ | 50 | 50 | 50 | 50 | 50 |
| | $N_{visit}$ | 3 | 3 | 3 | 3 | 3 |

### Model comparison

To examine ssMSOM performance, versus a typical analysis strategy for this type of data, we compared it to a series of single species models fit using maximum likelihood across all 30 spatial scales. These models are referred to as piecemeal GLMs throughout and are described by standard binomial GLM functions of the form:

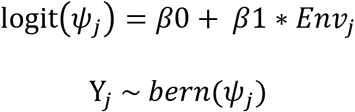

where *ψ_j_* is the naïve occupancy probability of the focal species at the focal spatial scale, and *Y_j_* is the naïve occupancy state (*i.e.,* whether a species was detected at a site across all site visits or not). To match the approach taken by empirical analyses species with observations at fewer than 10% of all sites were excluded from analysis (because data would presumably be insufficient for precise parameter estimates; *e.g.,* Desrochers *et al.* 2010; Zuckerberg *et al.* 2012). We then used AICc to choose the optimal spatial scale for each species in turn.

We focus on two core questions when evaluating the ssMSOM versus standard GLM approaches. First, does using an integrated community analysis provide more accurate estimates of the correct spatial scale (s) than a piecemeal approach? For species for which *βl* is close to 0, the spatial scale of response cannot be evaluated in the piecemeal GLMs because the scale of response is undefined if the species does not respond to the environment. For this set of analyses we therefore additionally excluded all species for which *βl* was not significantly different from 0 in the most likely GLM, as estimation of true spatial scale should be more accurate for the remaining species.

Second, even when estimating spatial scale is not the primary goal and is therefore considered a nuisance variable, does integrating over uncertainty regarding the correct spatial scale result in more accurate estimates of how species respond to the environment (*β*1) than a piecemeal approach? To answer this question we additionally compared parameter estimates with GLMs using the true spatial scale under which simulations were conducted. While for empirical data the true spatial scale is never known without error, using it here represents the ‘best-case scenario’ for community level analyses of landscape level responses to the environment.

To quantify the accuracy of the ssMSOM versus a piecemeal GLM approach, we calculated the root mean square error (RMSE) across the entire community of the family of parameter estimates from the true simulated value. With regards to ‘*s*’, we consider the posterior mode in the case of the ssMSOM, where as for the GLMs we consider the spatial scale that minimizes AICc for each species. To ensure the results are comparable in both approaches we calculate RMSE for each species in turn, even though in the case of the ssMSOM the parameter s is estimated for all species simultaneously, and is therefore identical for all species.

We also examine coverage probability of the posterior estimates (*i.e.,* the inverse of type I error). If models are behaving as expected, the 95% CIs of the parameter estimates should contain the true value 95% of the time. For ssMSOMs and GLMs the coverage probabilities for *βl* can be calculated directly from species-specific parameter estimates. Similarly coverage probabilities around ‘*s*’ for the ssMSOMs can be calculated using equal tail Bayesian credible intervals around the posterior of *s*. To calculate a value equivalent to coverage probabilities of the spatial scale in the case of the piecemeal GLMs we first calculated the AICc weight for all spatial scales for a given species, and then asked whether the true spatial scale was within the top 95% of the cumulative model weights.

### Model Fitting

Models were fit using JAGS through the R environment. Simulation code in R and JAGS model code is available in the supplement. For MCMC analyses diffuse priors were used throughout, with a flat prior placed on ‘*s*’.

## Results

### Inferring spatial scale

The posterior mode of spatial scales from the ssMSOM tended to accurately estimate the true spatial scale of response, and had relatively low error, typically off by less than 100m under the conditions simulated (Figures 2 and 3). In contrast, piecemeal GLMs failed to consistently recover the true spatial scale for the majority of species in the community, even when detection was perfect. The degree of error was lower when restricting analysis to only those species for which the lowest AICc GLM showed a significant relationship with the environmental gradient, though RMSE across the entire community was still >3X that of the ssMSOM (Figure 3a). Further, when detection itself varied along the environmental gradient at local scales in a species-specific manner, using a standard GLM approach resulted in error in estimating species response scales that is no better (and sometime worse) than guessing a scale at random (Figure 3a). Not surprisingly, error in estimating the scale of response within piecemeal GLMs was greatest for both species that were detected in a small number of sites, as well as species detected in the majority of sites (Figure 3B).

**Figure 2.**
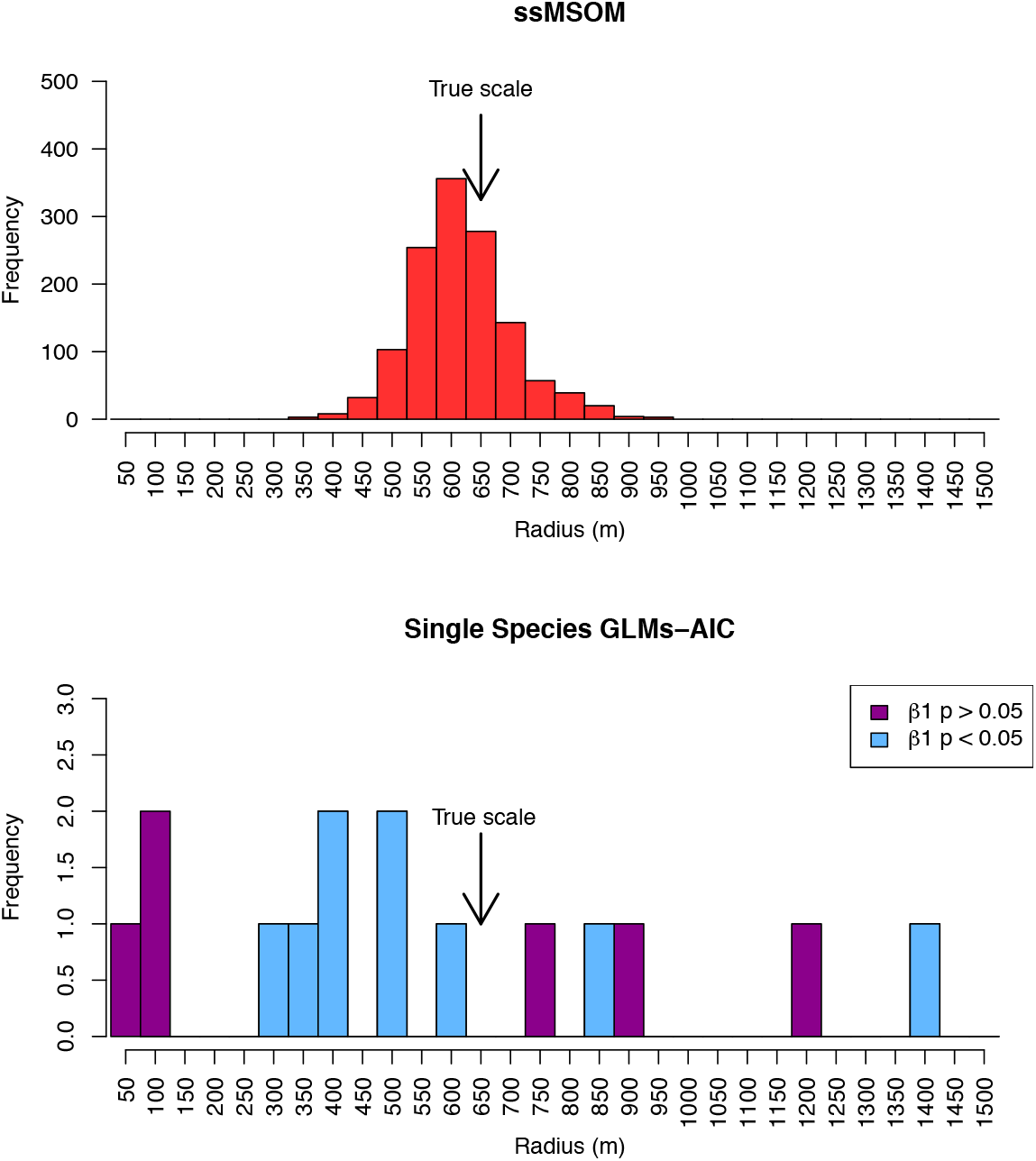
Example histograms of single species GLMs, and the ssMSOM. Example comes from low detection case (mean detection = ~0.25). Upper panel depicts posterior distribution of the spatial scale that describes species responses to the environment from the ssMSOM. Lower panel depicts the spatial scale that minimizes AICc for each single-species GLM, after removing one species that was observed in fewer than 10% of sites (i.e. 15 species remaining). Purple bars are species for which the response to the environment does not differ significantly from 0, at the ‘best’ spatial scale, whereas blue species have significantly positive or negative responses to the environment at the ‘best’ spatial scale.

**Figure 3.**
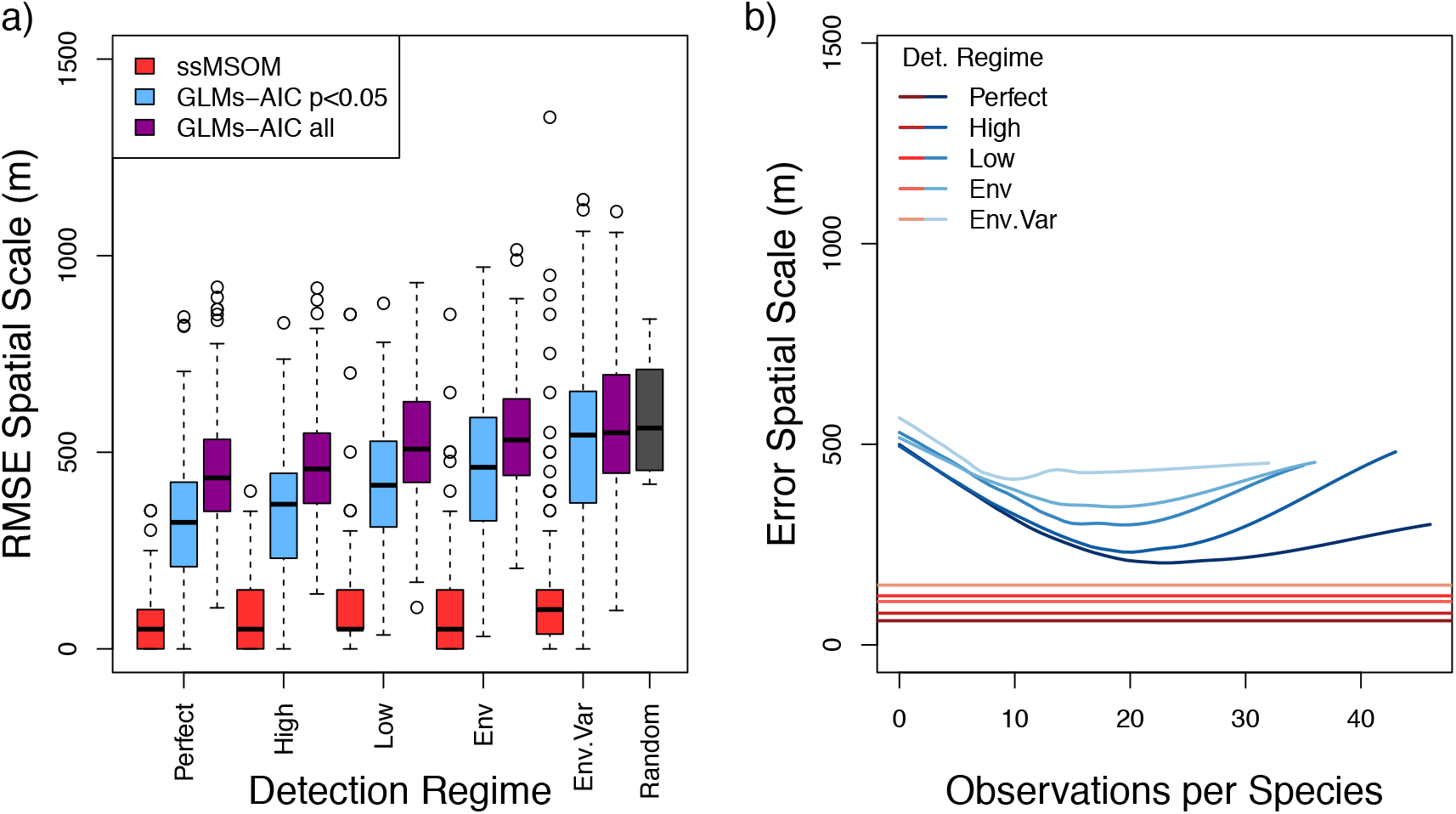
(a) General comparison of root mean square error for spatial scale across 120 simulations per detection regime, comparing posterior mode from ssMSOM, and AICc selected spatial scale for each species in single cases (splitting off only species with statistically significant responses to the environment [in blue], from all species [in purple]). ‘Random’ indicates the distribution of RMSEs that one would obtain if selecting spatial scales randomly along the uniform range from 50m to 1500m. (b) Blue lines in right panel depict lowess smoothers through all individual species’ error in spatial scale estimation from piecemeal GLMs (across all simulated communities), as a function of the number of individuals observed. Red lines show mean error across all ssMSOMs (all species in the community are assumed to respond at the same scale, so species’ level error is invariant to number of observations). Ability to estimate a species’ scale of response suffers when species are either too rare, or too common, when using piecemeal GLMs.

The ssMSOM demonstrated correct type I error rates when estimating s, regardless of detection regime. In contrast, piecemeal GLMs showed inflated type I error when estimating the true spatial scale, which was exacerbated as detection probability deviated from perfect (Figure 4). This behavior was further accentuated when excluding species that did not have a significant response to the environmental gradient at its most likely spatial scale, such that nearly 20% of all species assessments did not include the true spatial scale model in the top 95% Akaike weighted models under a low-detection regime with specific-specific variation in detectability by environment.

**Figure 4.**
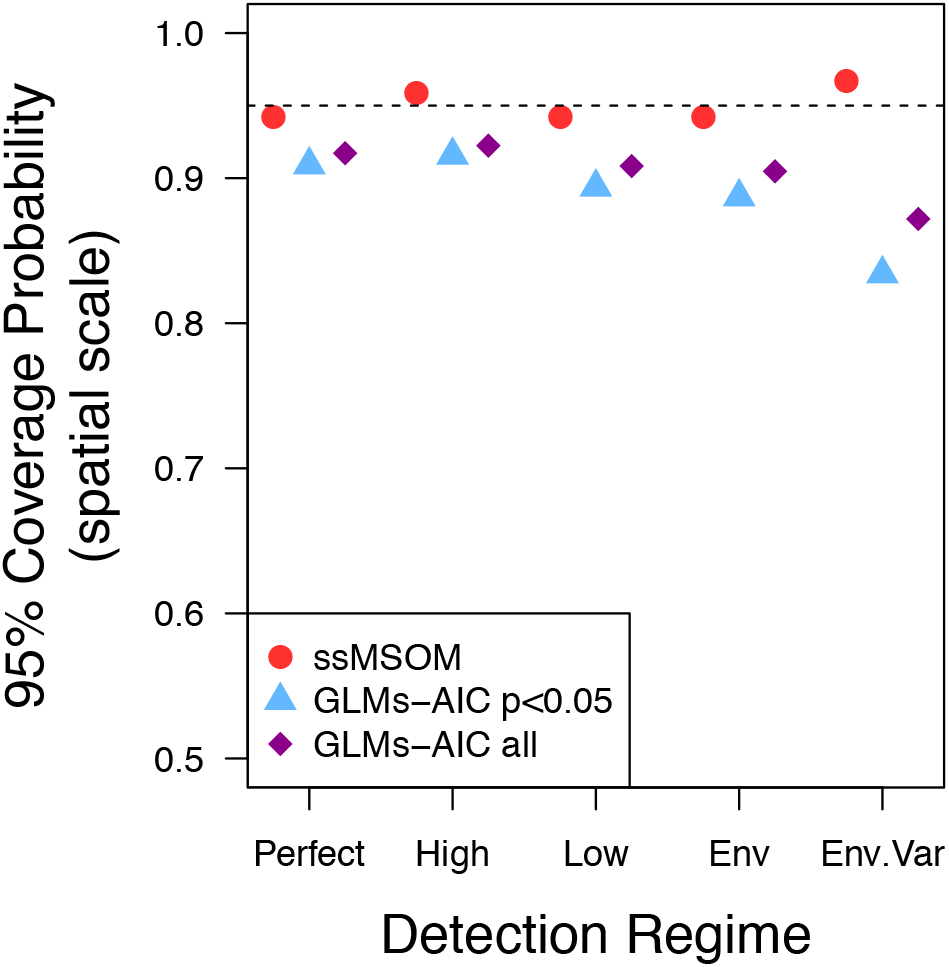
Quality of inference of true spatial scale declines for single species cases declines as detection becomes lower confounded with environment. For ssMSOM coverage probability indicates the proportion of simulations (n = 120 per regime) for which the true spatial scale was within the 95% CIs of the spatial scale posterior. For GLMs coverage probability indicates the proportion of species across simulations for which the true spatial scale was within the top 95% of Akaike weighted models.

### Estimating species responses to the environment

Estimates of species responses to the environment (*βl*) were more accurate in the ssMSOM than in piecemeal GLMs, even when GLMs were run using the true spatial scale (Figure 5). These patterns were not strongly affected by detection regime, though in general estimates are more accurate when detection probabilities are high. Type I error does however strongly shift with detection. If detection is perfect, and the true spatial scale in known *a priori*, then a piecemeal GLM approach performs as well as the ssMSOM (Figure 6). However, when the spatial scale must be inferred from the data GLMs generate falsely confident results, with the true values of species responses excluded from the 95% confidence intervals up to 30% of the time under some simulated conditions *(i.e.* 6X the nominal type I error rate). In contrast the ssMSOM possess 95% CIs that behave has expected, regardless of detection regime.

**Figure 5.**
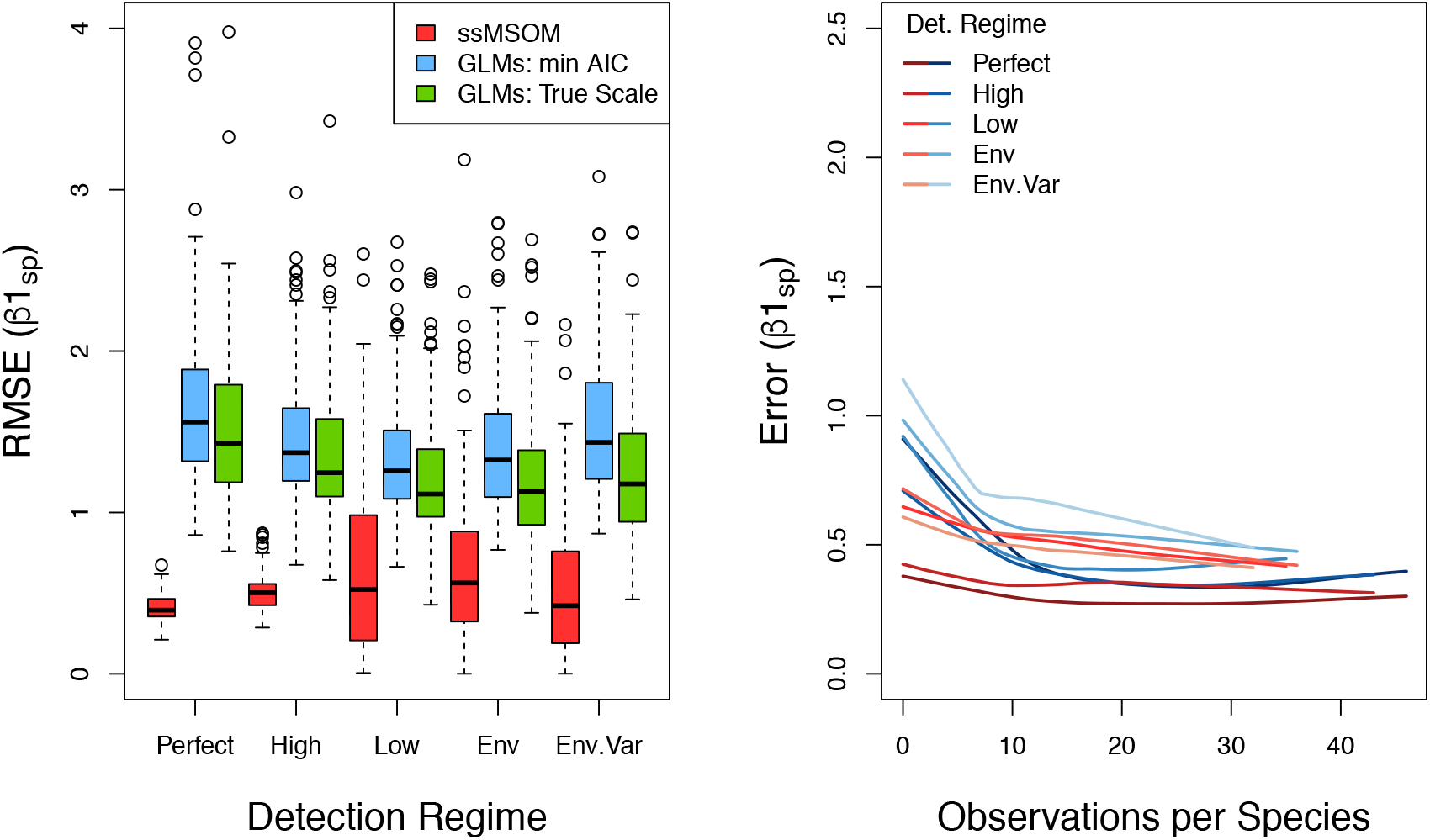
Accuracy of estimates of species responses to the environment (*βl*_sp_). Left panel depicts boxplots of community level RMSE of *βl*_sp_ values for each of 120 simulations per detection regime. Species observed in fewer than 10% of sites were removed from GLM estimates in left panel. Right panel depicts lowess smoothers through individual species’ error in *βl*_sp_ (across all simulated communities), as a function of the number of individuals observed. Here blue lines represent estimates from piecemeal GLMs (using min. AICc), whereas red lines represent estimates from ssMSOMs.

**Figure 6.**
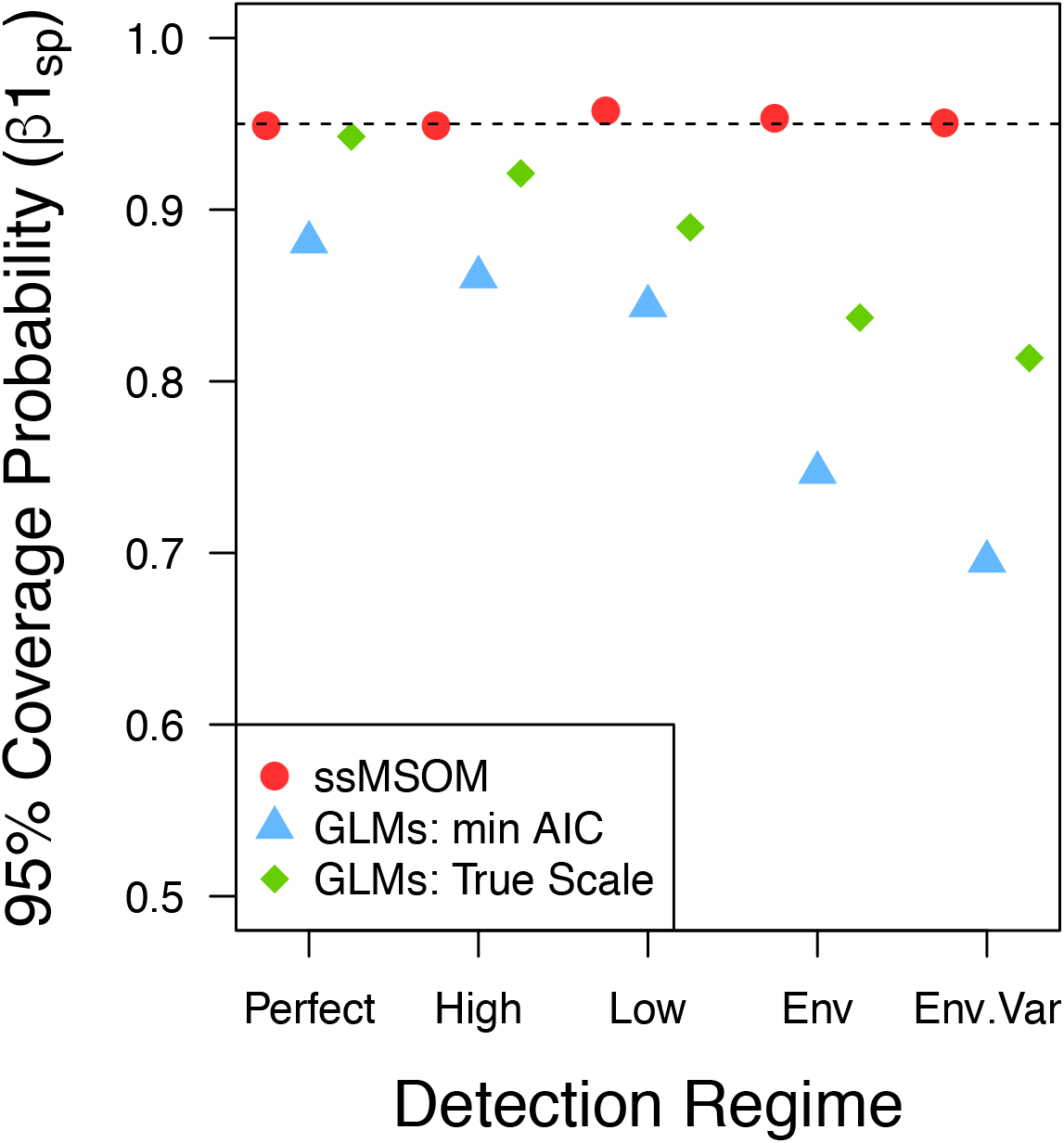
Coverage probabilities around true value of response to the environment (*βl*_sp_). Each point represents the proportion of species across all simulations for which the true value fell within the 95% CIs of the estimate. For GLMs all species observed in fewer than 10% of sites were excluded from the analysis. GLMs min AICc indicates each species’ GLM at the ‘best’ spatial scale. GLM True scale is the GLM from at the spatial scale that the responses were simulated at, regardless of whether this GLM possessed the lowest AICc.

## Discussion

Here we described and tested the statistical properties of the ssMSOM against the standard method for ascertaining species’ and communities’ scales of response. We find that internalizing scale selection into the model results in greater community wide accuracy for key parameter estimates, and reduces the probability of making incorrect inferences. The key strength of the ssMSOM is that it does not rely on setting the spatial scale *a priori.* Like the approach of Chandler & Hepinstall-Cymerman (2016), the ssMSOM avoids the problem of researchers selecting only a few scales to investigate, which are too narrow to include the true scale of response (Jackson & Fahrig 2015). This of course requires that researchers first extract landscape data from as broad a range of scales as possible, ideally in the finest increments possible. This allows spatial scale to be treated as nearly continuous, such that 95% CIs can be created, and inferences made as with any other continuous parameter in the model. When taking a flexible scale estimation approach it is essential to use as fine scale environmental data as possible. If environmental data are coarse with respect to the resolution at which species interact with the environment then the estimated spatial scale of response will be strongly upwardly biased and overall model performance will suffer (Mendenhall *et al.* 2011).

### Examples of empirical use

Two recent studies have demonstrated the power of using the scale selection routine from the ssMSOM (*i.e.,* indexing the *Env* matrix by scale) when analyzing empirical datasets. Frank et al. (in press) used a phylogenetic occupancy model (Frishkoff *et al.* 2017) with the internalized spatial scale selection method presented here, finding that bat responses to deforestation are strongly phylogenetically conserved. Similarly, Karp et al (in review) used Bayesian spatial scale selection embedded within an *N*-mixture model to examine how bird communities responded to habitat conversion while accounting for imperfect detection in order to understand how *β*-diversity was structured along land-use and climate gradients. In both cases spatial scale selection strongly supported deforestation affecting the communities at fairly small spatial scales. While in both cases scales at over a kilometer away from focal sites were queried, for bats the posterior distribution peaked at 50m, and excluded all scales above 100m, while for birds scales below 300m were favored. Because of the ssMSOM framework, these studies were able to analyze both common and rare species. Had these studies relied on individual GLMs (or species-by-species occupancy models) the uncertainty around the scale of response would likely have been extremely high, and un-estimatable for the majority of rare species. This was particularly important in the case of Neotropical bats for which rare and hard to detect species tended to be found in natural forests. Indeed, if imperfect detection were not taken into account species richness would have appeared to have been unaffected by forest loss, when in fact it declined sharply (Frank *et al. In press).* These early examples of embedding spatial scale selection into hierarchical models highlight the broad applicability of the method. The ssMSOM approach is easily extended to abundance models (i.e., *N*-mixture or recapture models), or indeed any Bayesian implementation of multispecies models with or without detection for which the true spatial scale of response is unknown could benefit from the general approach.

### Assumptions, limitations, and future directions

Critically, for the simulations presented (and within the ssMSOM itself) there is the assumption that a single, true spatial scale exists at which all species respond to the environment. This may or may not be true for a given assemblage in nature. Empirical studies have shown that different species respond to different spatial scales (*e.g.*, Chambers *et al.* 2016), and theoretical approaches suggest that some species traits may modulate the scale of response (Jackson & Fahrig 2012). However, we show in our simulations that empirical analyses conducted on a species by species basis (as past studies have been) are often unable to recover the true spatial scale at which species respond, and show high heterogeneity in the scale of response even if all species are simulated to respond at the same spatial scale. While many species likely do respond at different scales, this finding casts some doubt on the specific estimates of scales of response presented in past empirical studies. The high degree of inaccuracy inherent in the piecemeal GLM approach may be partially responsible for the lack of correlation between empirically estimated scales of response, and species traits thought to modulate these scales (Jackson & Fahrig 2015).

Future development of the ssMSOM and similar community wide approaches should be able to relax the assumption that all species have the same scale of response, although doing so may diminish the ability to precisely estimate response scales for rare species. One path would be to estimate spatial scale separately for two or more groups of species, delimited based on natural history knowledge, functional guild placement, or other *a priori* expectations (Pacifici *et al.* 2014). An *a priori* grouping based approach, however, is at best an imperfect solution. Ideally individual species’ scales of response would be allowed to vary from one another—using random effect structures within the scale selection component of the model would be one logical way to do so. Allowing random variation among species could additionally allow species’ level covariates to affect the scale of response, thereby facilitating testing the hypothesis that some species’ traits correlate with scale of response that both maintains high power, and is minimally afflicted by type I error.

In real communities, species’ responses to the broader landscape might be predicated on conditions at the local scale (*i.e.,* an interaction between local and landscape scales). For example, a farm-land bird species might benefit from landscape level tree cover when the local habitat is agriculture, but might only exist in forest when there are low amounts of landscape level tree cover because it uses forest edge habitat. Allowing interaction terms between landscape and local effects (e.g., forest cover as estimated within a point count radius) will allow these types of species interactions with the environment to be tested (Matthiopoulos *et al.* 2011; Paton & Matthiopoulos 2016).

Chandler and Hepinstall-Cyberman (2016) pointed out that the step function used to calculate proportion of focal habitat within a given radius has no theoretical basis, and instead favor a Gaussian weighting function. This approach could be easily implemented with the Bayesian framework presented here, by indexing the *Env* matrix based on the output of the weighting function over incremental changes in its key parameter. While, alternatives to the commonly used step functions are certainly appealing on theoretical grounds, at least one study that examined Gaussian weighting versus a step function radius method found that models performed roughly equivalently (Timm *et al.* 2016).

### Conclusion

Humans are altering landscapes across the globe, such that the remaining extent of natural habitats are often much diminished and severely fragmented (Haddad *et al.* 2015). Such complex, heterogeneous landscapes challenge ecologists’ abilities to discern the underlying environmental drivers of community composition. Yet achieving successful conservation strategies in these landscapes requires simultaneously describing and predicting how these spatially heterogeneous environments affect not just individual species, but entire communities. Internalizing spatial scale selection within community models offers one approach to uncover the environmental drivers behind such community change while accommodating the unavoidable uncertainty in the ‘true’ scale of species’ responses. The ssMSOM possess high accuracy and correct type I error rates when both identifying the spatial scale of response, and the direction and magnitude with which individual species respond to environmental gradients. This approach represents a promising path forward for understanding the ecological drivers of community composition, and the consequences of ongoing environmental change.

## Acknowledgements

We would like to thank D. S. Karp and A. Echeverri for use of fine-scale forest cover data from northwestern Costa Rica. This study was supported by a University of Toronto Ecology and Evolutionary Biology Postdoctoral Fellowship to LOF and NSERC Discovery Grants to DLM and M-JF.

### Data Accessibility

R scripts for simulating communities and running ssMSOMs are available in the supplemental material.

